# Doppler Slicing for Ultrasound Super-Resolution Without Contrast Agents

**DOI:** 10.1101/2021.11.19.469083

**Authors:** Avinoam Bar-Zion, Oren Solomon, Claire Rabut, David Maresca, Yonina C. Eldar, Mikhail G. Shapiro

## Abstract

Much of the information needed for diagnosis and treatment monitoring of diseases like cancer and cardiovascular disease is found at scales below the resolution limit of classic ultrasound imaging. Recently introduced vascular super-localization methods provide more than a ten-fold improvement in spatial resolution by precisely estimating the positions of microbubble contrast agents. However, most vascular ultrasound scans are currently performed without contrast agents due to the associated cost, training, and post-scan monitoring. Here we show that super-resolution ultrasound imaging of dense vascular structures can be achieved using the natural contrast of flowing blood cells. Instead of relying on separable targets, we used Fourier-based decomposition to separate signals arising from the different scales of vascular structures while removing speckle noise using multi-ensemble processing. This approach enabled the use of compressed sensing for super-resolution imaging of the underlying vascular structures, improving resolution by a factor of four. Reconstruction of ultrafast mouse brain scans revealed details that could not be resolved in regular Doppler images, agreeing closely with bubble-based super-localization microscopy of the same fields of view. By combining multi-ensemble Doppler acquisitions with narrowband Fourier decomposition and computational super-resolution imaging, this approach opens new opportunities for affordable and scalable super-resolution ultrasound imaging.

## I. Introduction

Ultrasound is one of the most widely used imaging modalities due to its portability, cost-efficiency, and high temporal resolution. However, only the recent introduction of ultrasound super-localization microscopy (ULM) enabled ultrasound to resolve microvascular structures deep within the tissue [1]–[4]. The order of magnitude improvement in spatial resolution and high-contrast images produced by super-localization microscopy quickly found applications in neuroimaging and cancer research [5]–[7]. However, despite the continuous spread of contrast-enhanced ultrasound imaging, most vascular ultrasound scans are still performed without injecting contrast agents. Instead, these vascular scans rely on the natural acoustic contrast of our blood – the scattering from flowing blood cells [8]. The reliance on Doppler scans is especially widespread in point-of-care scans, remote radiological readouts, and clinics in developing countries. Therefore, this work explores the feasibility of performing super-resolution ultrasound imaging of dense vascular structures without ultrasound contrast agent injections.

Ultrasound super-localization methods surpass the acoustic diffraction limit by precisely estimating and aggregating the locations of separable microbubbles over many consecutive frames [1], [2]. Several approaches for ultrasound super-resolution imaging have relaxed the requirement for target separability, enabling higher contrast agent concentrations and shorter acquisition times. Instead of relying on separable targets, different properties of contrast-enhanced ultrasound signals were used, including the temporal fluctuations of blood flow signals [9], the sparsity of the underlying vasculature [10], [11], and the flow trajectories of the bubbles [12]. Initially, these frameworks relied on iterative algorithms for sparse representation and compressed sensing [10]. The subsequent adaptation of deep learning further improved the resolution by capturing additional underlying structures in these signals [11], [13], [14]. These advancements contributed to the development of fast microbubble super-resolution imaging methods [15], [16].

Direct super-resolution imaging of the microvasculature without contrast agents is more challenging because contrast-free ultrasound scans have a lower signal-to-noise ratio (SNR) and less inherent sparsity compared to contrast-enhanced scans. Typically, in contrast-enhanced scans, only a subset of the vessels contains microbubbles at any given time [17]. However, even dense vascular structures include vessels with different flow directions and velocities that could facilitate the decomposition of non-contrast vascular signals to a series of sparser components. While directional Doppler display is commonly used [18], a much finer separation of vascular signals is needed for increasing the sparsity of non-contrast vascular scans.

A similar challenge of increasing sparsity in narrowband radar imaging was tackled using a method called Doppler focusing [19]. Doppler focusing decomposes the data to single Fourier bands according to the different velocities of the imaged targets. Such narrowband processing in which the Doppler spectrum is broken down to hundreds of bands is distinct from previous wideband ultrasound imaging approaches that cover the entire spectrum with two or four filters [10]. However, unlike radar, in contrast-free ultrasound imaging, there are many scatterers per resolution cell. The resulting constructive and destructive interference gives rise to the dominant multiplicative speckle patterns that obscure the vascular information in single Fourier bands (**Fig. 1**). To remove this time-dependent speckle pattern, we integrated the vascular information in each band over tens to hundreds of Doppler acquisitions. As presented in this paper, this approach for multi-acquisition narrowband processing, termed Doppler slicing, reveals high SNR vascular details, crucial for the successful application of sparse representation algorithms [20], [21]. Thus, Doppler slicing could open new opportunities for super-resolution ultrasound imaging enabling sub-diffraction ultrasound imaging without contrast agent injections.

**Figure 1:**
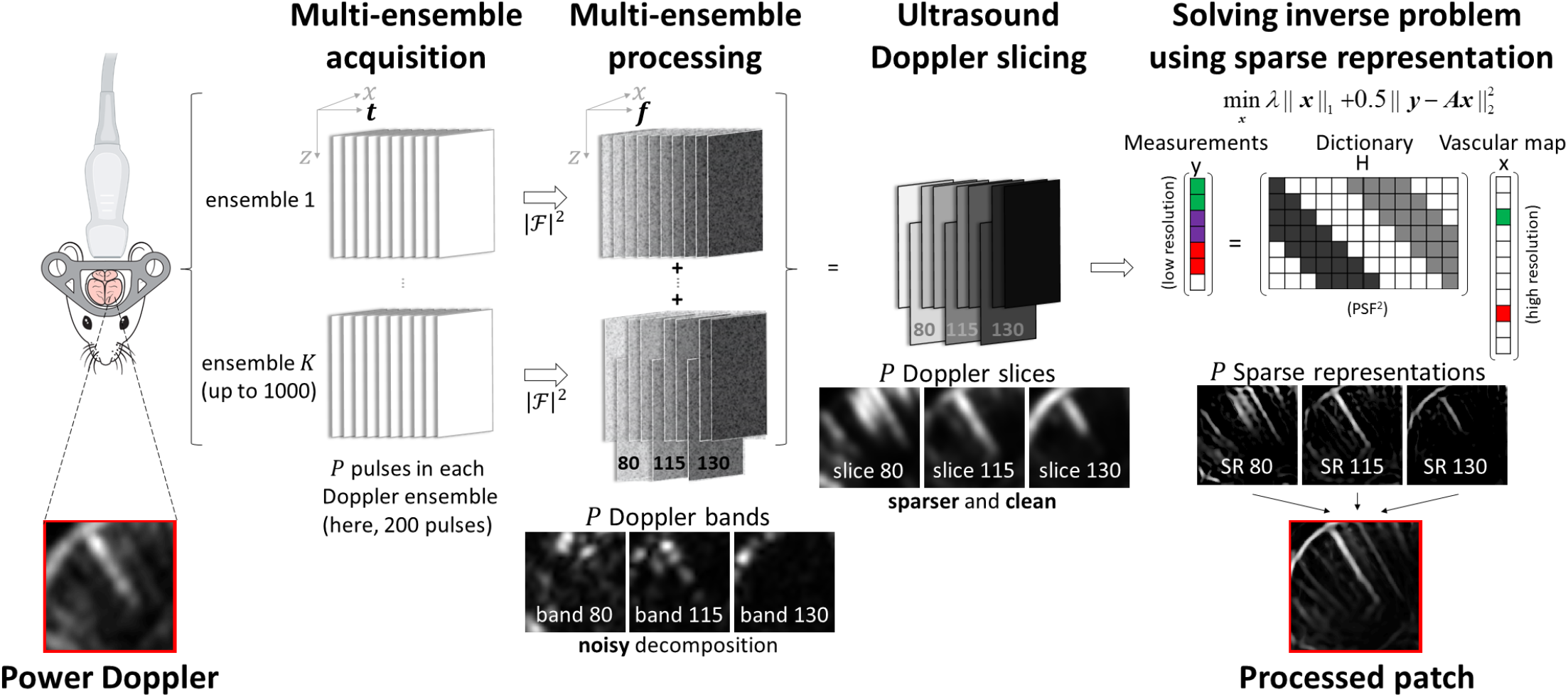
Ultrasound Doppler slicing results in sparser and clean vascular decompositions. Ultrasound Doppler ensembles are decomposed into several Doppler bands, each containing a sub-set of the imaged vascular structure. Subsequently, the vascular information is integrated over many consecutive ensembles removing the strong speckle-noise patterns from the vascular maps. Doppler slicing facilitates super-resolution vascular imaging by producing a series of sparser vascular images with a high signal-to-noise ratio and dynamic range. Images are 1.6 mm wide. The figure was created with BioRender.com.

The rest of the paper is organized as follows: A parametric model for non-contrast ultrafast ultrasound hemodynamic signals, including Doppler information, is discussed in Section Section III shows how Doppler focusing can be adapted to ultrasound imaging and discusses the unique challenges in performing such decomposition. The complete processing scheme combining Doppler decomposition and sparse representation is introduced in Section IV. In Sections V and VI, we show how the proposed approach results in super-resolution ultrasound imaging of intricate vascular structures without contrast agents. We demonstrate the feasibility of our method on neural mouse scans and show up to four-fold improvement in spatial resolution. Our results are analyzed and discussed in Section VII, which concludes the manuscript.

Throughout the paper, *x* denotes a scalar, ***x*** a vector, and ***X*** a matrix. The size of a matrix ***A*** is denoted by *MXM*, with ***A*** having *M* rows and *M* columns. Square brackets [·] represent discrete-time signals, and round brackets (·) indicate continuous-time signals. The *k*th discrete Fourier transform coefficient of *x*[*p*], *p* = 1, …, *P* is denoted using capital letters and the index *k, M*[*k*], *k* = 1, …, *P*. The notation || · ||_*p*_ indicates the standard *p*-norm.

## II. Model for Non-contrast Ultrafast Ultrasound Hemodynamic Imaging

In the past decade, ultrasound plane wave imaging has gained increasing popularity, providing an order of magnitude higher frame rate and dramatically better vascular detection than classic focused beam imaging [22]. Following each plane wave pulse, the echoes from the imaged tissue are received by all the transducer elements. Then, received data is focused computationally via beamforming, producing a spatial map of ultrasound reflections [23]. For example, classic delay and sum beamforming is performed by applying the proper time delays to each data channel before summing them [23]. The beamforming process defines a spatial grid on which the signal is sampled with spacing [Δ_*xL*_, Δ_*ZL*_] in the lateral and axial directions, respectively.

In pulse-echo Doppler measurements, *P* equally spaced ultrasound scans are acquired for each blood flow measurement. This packet of pulse-echo measurements is known as a Doppler ensemble, and the entire span of the Doppler ensemble is called the coherent processing interval (CPI) [24]. The blood-related signal is contaminated by the tissue clutter *c* and an additive thermal noise component *w*. Extending on the model for contrast-enhanced ultrasound signals in [10], after beamforming and demodulation, the *IQ* signal *f* of hemodynamic ultrasound scans can be written as:

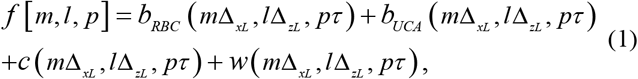

where, *b*_*RBC*_ is the signal coming from the blood cells and *b*_*UCA*_ is the signal originating from contrast agents flowing in the bloodstream. Here, *m* is the index of the grid in the lateral direction, *l* is the index of the grid in the axial direction, *m, l* ∈ {1, …, *M*}, *p* is the pulse number within the Doppler ensemble and *τ* is the pulse-repetition interval. In contrast to previous studies [9], [10], in this work, ultrasound contrast agents are not injected and therefore *b*_*UCA*_ = 0. The only blood-related signal *b*_*RBC*_ is composed of the echoes scattered from numerous blood cells. This source of signal was neglected in previous related works since it is significantly weaker than echoes coming from microbubbles. The blood signal is frequently estimated from ultrafast ultrasound scans via singular value decomposition (SVD) filtering, removing the highly coherent tissue-related signal from the low coherence blood flow signal [25]. For ease of notation, from here on, *b* represents the blood flow signal *b*_*RBC*_ estimated from *f*.

Hemodynamic ultrasound measurements use many repeated Doppler acquisitions to quantify blood flow changes over the cardiac cycle or in response to external stimulation. In functional ultrasound (fUS) scans, Doppler-based measurements of the deviation from the baseline flow level are used to infer brain activity [18]. While each ensemble is acquired using an equally spaced synchronous stream of pulses, the different ensembles are frequently asynchronous due to hardware implementation considerations [18]. In each of many repeated Doppler measurements, the flow level in each pixel is estimated by calculating the Doppler signal’s power over the CPI. This method of measuring and presenting Doppler signals, called power Doppler display, is one of the common ways of depicting flow in vascular ultrasound scans. In fUS, each Doppler ensemble is processed separately, and power Doppler changes in each pixel are tracked over time [26]. In contrast, this work investigated the use of up to 1000 ensembles and processed them together to gain more precise information about the underlying vascular structures, extending the discussion beyond classic ultrasound Doppler imaging. Notably, the pulsing parameters of typical plane wave acquisitions allow these 1000 ensembles to be acquired in under three and a half minutes.

The signal coming from each blood cell can be presented as a convolution between its reflectivity function and the system’s point speared function (PSF). The PSF of the ultrasound system *h*[*m, l*], varies with the location in both the axial and lateral direction. However, it can be assumed to be piece-wise constant when processing small image regions (or patches), similar to the approach taken in [27] for focused-beam imaging. It is important to note that blood cell scattering can be assumed to be linear in relation to the pressure of the incident wave. In comparison, the echoes emitted from each microbubble in contrast-enhanced ultrasound imaging depend on the pressure wave in a non-linear way, determined by the bubble size and other shell properties that change from bubble to bubble [28]. en looking at each image patch, blood cells’ scattering is adequately represented by the same PSF. For each Doppler ensemble *k*, the blood-related signal *b*_*k*_ can be expressed as a superposition of the scattering from all the cells:

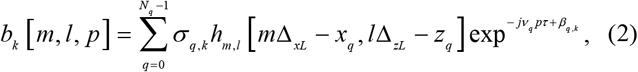

where *M*_*q*_ is the number of scatterers, and *x*_*q*_, *z*_*q*_ are the position of each scatterer in the lateral and axial directions, respectively. The scattering of blood cell *q, σ*_*q,k*_ is equal to 0 in ensembles for which the blood cell is outside the imaging plane. Here, *v*_*q*_ is the Doppler angular frequency, and *β*_*q,k*_ is a random constant phase determined by the initial location of the scatterer relative to the transducer. Each Doppler frequency is related to a specific axial velocity *v*_*z,q*_, by the following relation,

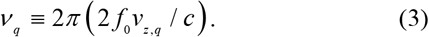

Here, *c* is the velocity of sound in the medium and *f*_0_ is the central frequency of the ultrasound pulse. The flow level and the velocity of blood flow are the main parameters estimated by hemodynamic ultrasound imaging.

Several assumptions can be made regarding the measured microvascular Doppler signals, simplifying the model and subsequent analysis. These assumptions are similar to those used in radar imaging [19]:

1. “Far targets”: the distance between the transducer and the scatterer is large compared to the distance traveled by the scatterer during the CPI (0.2 s in our case). This assumption allows us to treat *σ*_*q,k*_ as a constant for each ensemble.
2. “Slow targets”: the target velocity is small enough to assume that *x*_*q*_ and *z*_*q*_ are constant throughout the CPI.
3. “Small acceleration”: the target velocity remains approximately constant throughout the ensemble, allowing for constant *v*_*q*_.
4. “No time ambiguity”: The time delay between consecutive imaging pulses is such that the image target is within the desired imaging depth: 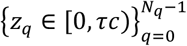
5. “No Doppler ambiguity”: the measured Doppler frequencies are within the unambiguous frequency region: 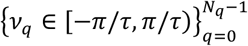

The described signal model and related assumptions will guide us in defining the Doppler slicing operation and sparse representation processing described in the following sections.

## III. Ultrasound Multi-acquisition Doppler Slicing

Non-contrast enhanced Doppler scans contain a much higher density of scatterers than contrast-enhanced ultrasound acquisitions [17]. Unless scanning poorly perfused tissues like the cores of certain tumors, a single volume cell frequently contains blood cells flowing in several vessels with different Doppler angular frequencies *v*_*q*_. As we will see in the following sections, the lower SNR and higher density of non-contrast ultrasound Doppler measurements limit the use of sparsity-exploiting methods when processing these scans. Therefore, to highlight the underlying spatial structure, we decompose the received blood signals based on angular frequency, a process that results in a sparser vascular network in each sub-dataset.

Inspired by the concept of radar Doppler focusing [19], we define the following Doppler slicing function that decomposes the received blood signal into P frequency bands (Fig. 1):

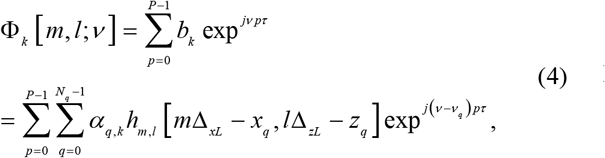

using (2) and representing the complex amplitude of each scatterer as 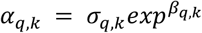. Denoting 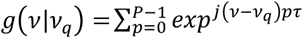 we get:

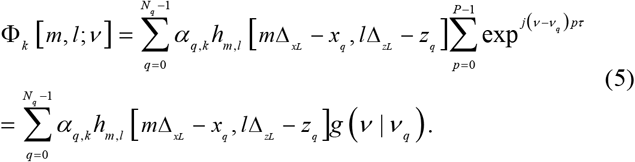

For any Doppler band *v*, echoes originating from blood cells flowing in a band of width 2*π*/(*Pτ*) around it will be integrated coherently as the phase between *v* and *v*_*q*_ is close to zero [19]. This range around *v* will be considered as the *v* “Focus zone”. Therefore, for a single blood cell within the focus zone, we can write:

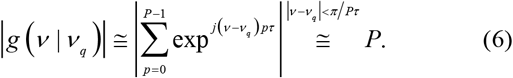

Vascular details within the *v* “Focus zone” will be preserved while targets outside the focus zone will approximately cancel out. As a result, complying with the Fourier transform definition, larger vessels with higher blood flow velocities will be included in the middle bands while microvascular details will be concentrated in the first and last few Doppler bands.

To study the finer details of the vasculature, we will define a high-resolution spatial grid. We introduce a high-resolution grid with spacing [Δ_*xH*_, Δ_*zH*_], such that the location of the scatterers can be attributed to one of these pixels and [*x*_*q*_, *z*_*q*_] ≅ [*i*_*x*_ [*q*]Δ_*xH*_, *i*_*z*_ [*q*]Δ_*zH*_] for some *i*_*x*_, *i*_*z*_ ∈ {0, …, *N*− 1}, similar to [10]. Thus, the Doppler slicing function can now be approximated by:

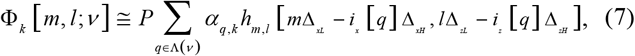

where Λ(*v*) = {*l*: *| v* − *v*_*q*_ *|*< *π*/*Pτ*. From here on, we will refer to [Δ*xL*, Δ*zL*] as the low-resolution grid. Assuming that Δ_*xL*_ = *D* Δ_*xH*_ and Δ_*zL*_= *D* Δ_*zH*_ for some *D* ≥ 1, it holds that *N* = *DM*. Substituting Δ_*xL*_ = *D* Δ_*xH*_, Δ_*zL*_= *D* Δ_*zH*_ into (7) we have

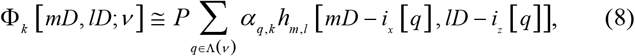

where Δ_*xL*_, Δ_*zL*_, Δ_*xH*_ and Δ_*zH*_ are omitted for convenience. Studying the blood cells’ echoes in each Doppler band on this high-resolution grid will help infer the underlying vasculature structure.

In contrast to super-localization microscopy, where single bubbles can be isolated, the scattering of a single blood cell is too low to detect, and their density and enormous number make them inconceivable to resolve. Therefore, we focus on separating blood vessels and not the blood cells they contain. Consequently, we sum all the complex amplitudes of the scatterers that are included in each high-resolution pixel [*i*_*x*_, *i*_*Z*_] and define the complex amplitude of the blood signal in that pixel as 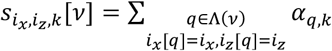 :

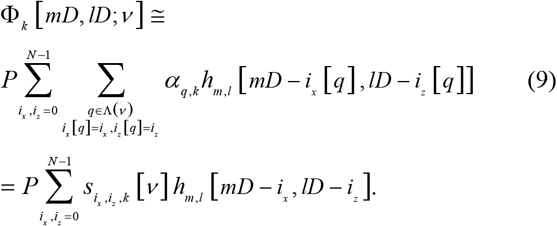

The blood signal in each pixel, 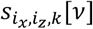, is governed by the complex amplitudes of the blood cells’ echoes and, therefore, by the flow level in each pixel. For example, in pixels that do not include any blood vessels at the inquired Doppler band, 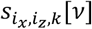 will be equal to zero. However, the magnitude of 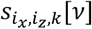 will change between ensembles according to the exact number and locations of the blood cells. These randomly varying complex amplitudes are the source of the speckle noise characteristic of vascular ultrasound data [29].

To facilitate the analysis of the randomly changing blood flow signals, we make the following additional assumptions:

6. “Statistical independence”: It is assumed that the contributions of the flow in different blood vessels to the overall measured signal are statistically independent. Therefore, 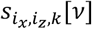 in different vessels can be assumed to be independent over the K repeated Doppler measurements and 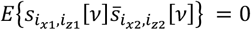 for *i*_*x*1_ ≠ *i*_*x*2_ and *i*_*z*1_ ≠ *i*_*z*2_.
7. “Motionless vasculature”: While the blood cells are constantly moving, the locations of the blood vessels do not change during this short acquisition time. The assumption of motionless vasculature can be relaxed using proper motion compensation [30], [31].
8. “Steady flow”: Without applying external stimulation, we can assume that the baseline blood flow level doesn’t change between repeated Doppler ensembles. Therefore, as a function of k, 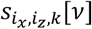 are independent and identically distributed random variables. This assumption does not hold in contrast-enhanced ultrasound since bolus injections result in quick wash-in and long clearance phases.

These assumptions enable us to extend the processing beyond classic power Doppler imaging and integrate the information on the vascular structure over several ensembles while removing the random speckle-noise patterns.

The presence of many scatterers in each resolution cell and the significant visual artifact of speckle-noise (**Fig. 1**) are not central in radar Doppler focusing. Since this speckle pattern changes between acquisitions, the vascular map in each band was estimated by averaging the squared magnitude of the Doppler slicing function in each pixel over the K ensembles:

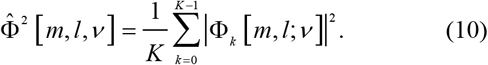

The expected value of the squared Doppler slicing function is equal to:

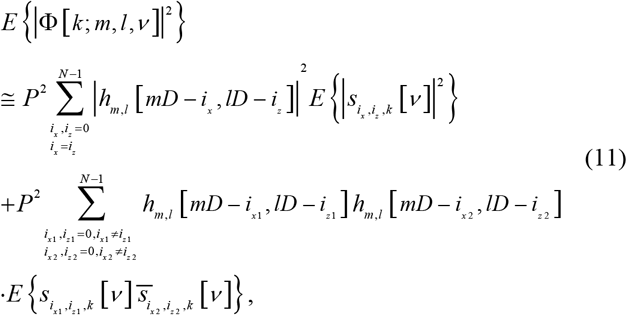

where, [*i*_*x*1_, *i*_*z*1_] and [*i*_*x*2_, *i*_*z*2_] are the coordinates of high-resolution pixels located along the same streamline. Blood cells flowing independently in different vessels produce only the first squared absolute-valued PSF term in equation (11) without cross-terms. Therefore, the cross-term in (11) can be neglected in our analysis, as it does not affect the separation of neighboring blood vessels, similar to [10]. Neglecting the second term in (11), we get:

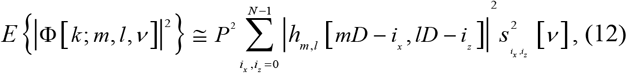

where 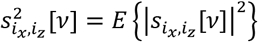 is the expected value of the blood fluctuations’ magnitude in the high-resolution pixel [*i*_*x*_, *i*_*Z*_]. Being narrower than the original PSF, the squared PSF represents an improved separation between the vessels. By combining multi-ensemble processing with Doppler slicing decomposition, we get a series of clear vascular maps, separated according to blood flow hierarchy (**Fig. 1**).

In addition to the decomposition of vascular structures, Doppler slicing holds two characteristics important for estimating the vascular anatomy while exploiting its underlying sparsity: improved signal-to-noise ratio (SNR) and dynamic range (DR). Radar Doppler slicing spreads the thermal noise over all the Doppler bands, resulting in an SNR improvement by a factor of *P* [19]. In ultrasound scans, each blood vessel includes a distribution of blood-flow velocities. Small vessels are captured by only a few Doppler components and will benefit the most from improved SNR following Doppler slicing since they are closer to the noise floor. In comparison, the ultrasound signals measured from large blood vessels are spread over several Doppler bands. However, these large blood vessels are clearly depicted in each Doppler band. In addition, since large blood vessels have stronger ultrasound signals, separating large blood vessels from smaller ones using Doppler slicing also improves the dynamic range in the slower blood flow bands. Together, the increased sparsity and improved SNR and DR result in the ability to detect and resolve smaller vessels from scans of dense vascular structures.

## IV. Sparse Representation of Ultrasound Doppler Slices

After defining the Doppler slicing operation and the high-resolution grid, we can now describe the sparse representation processing scheme that results in super-resolution vascular imaging. Following Doppler slicing, we assume that on the high-resolution grid, the underlying vasculature in each band is sparser, lending itself to the formulation of a sparse recovery problem [20], [21]. Instead of neglecting the information given by the Doppler velocities, by applying Doppler slicing, we look at the problem of locating only the blood vessels that belong to a specific Doppler band, implicitly estimating the range of Doppler frequencies in each blood vessel. This problem can be solved using iterative algorithms, such as in this work, or machine learning approaches that can uncover additional structures within the dataset.

We aim at reconstructing an *NXN* super-resolved image of the vasculature, which is *D*^2^ times larger from the initial *MXM* IQ matrix. This *NXN* matrix, 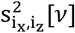 represents the variance of the blood-related echoes in pixel [*i*_*x*_, *i*_*Z*_]. This variance 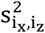 will be equal to zero in every pixel that does not contain blood vessels. By estimating the map of 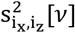 values, a high-resolution estimation of the vascular structure can be achieved.

Following a similar line of computation to that was presented in [32] and used in [10], we consider (12) in the discrete Fourier domain, which leads to efficient numerical estimation of the high-resolution image. While the Doppler slicing step is based on temporal Fourier decomposition, this sparse representation algorithm includes spatial Fourier transform. We denote the 2D discrete Fourier transform of 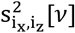 as *S*_2_[*k*_*m*_, *k*_*l*_, *v*], where *k*_*m*_, *k*_*l*_ are *MXM* spatial frequencies. Performing an *MXM* 2D DFT on (12) yields

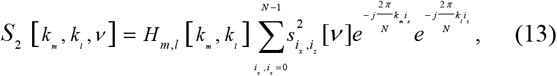

where *H*_*m,l*_ [*k*_*m*_, *k*_*l*_] is the *MXM* 2D DFT of the squared, absolute value PSF |*h*_*m,l*_ (*xD, yD*)|^2^. Next, we rewrite (13) using matrix-vector notation. To simplify the equation, we perform column-wise stacking of *S*_2_[*k*_*m*_, *k*_*l*_, *v*] and write the result as an *M*^2^ long vector ***y*** [*v*] = *vec*{*S*_2_[*k*_*m*_, *k*_*l*_, *v*]}. Similarly, we vectorize the *MXM* blood-flow statistics 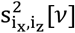 on the high-resolution grid and write the result as an *N*^2^ long vector 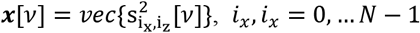. Thus, ***x***[*v*] represents the underlying vasculature in band *v* that we wish to recover on the high-resolution grid. Rewriting (13) in matrix-vector form, we get:

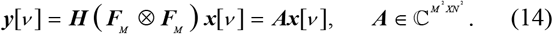

Here, ***A*** = **H** (***F***_*M*_⨂***F***_*M*_), ***H*** is an *M*^2^*X M*^2^ diagonal matrix with diagonal elements {*H*[0,0], …, *H*[*M* − 1, *M* − 1]}, ⨂ denotes the Kronecker product and ***F***_*M*_ represents a partial *MXN* DFT matrix, calculated by taking the rows corresponding to the lowest *M* frequency components of a full *NXN* DFT matrix.

Exploiting sparsity enables reconstruction of the underlying vascular structure at sub-diffraction resolution. For each image patch, we estimate ***x***[*v*] in (14) by solving the following optimization problem, assuming that ***x*** is sparse:

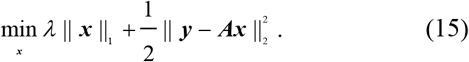

Here, *λ* ≥ 0 is a regularization parameter, and *v* is omitted for the sake of clarity. ***x*** represents the variance of the blood flow signal fluctuations, which is a non-negative quantity, therefore the constraint ***x*** ≥ 0 is added. Many existing algorithms aim at solving (15). As in [10], we used the FISTA [33] algorithm, which is known to achieve the fastest possible (worst-case) convergence rate for a first-order method, as described by Nesterov [33]. A detailed description of this implementation can be found in [10].

The application of sparse representation to each Doppler band’s blood flow variance map results in a series of super-resolved vascular images. A super-resolved map of the full vasculature is then reconstructed by integrating over the *P* different bands and combining all the image patches:

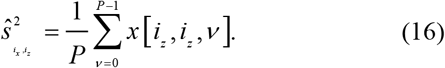

The following sections show that sparsity-based reconstruction is significantly improved by applying Doppler slicing to the low-resolution measurements.

## V. Materials and Methods

The proposed framework was validated using numerical simulations and tested on *in vivo* ultrasound scans of mouse brains. The cranial vascular scans were decomposed using Doppler slicing to characterize this method’s performance when applied to dense vascular structures. The decomposed datasets were then used to study the ability of Doppler slicing to facilitate super-resolution reconstruction when combined with sparse representation processing. The blood flow simulations helped us interpret these results by evaluating the performance of the sparse representation algorithm under different levels of SNR and imperfect PSF estimations.

### A. Numerical simulations

The custom-made numerical simulations used in this study included two blood flow geometries. In the first one, two parallel blood vessels with blood flowing in the same direction were positioned next to each other to measure this technique’s resolution improvement and vessel separation capabilities. These experiments were designed to test performance of sparse representation algorithms when applied to blood flow data without contrast agents. Therefore, unlike previous studies [9], [10], these simulations included more than ten scatterers per resolution cell, resulting in conditions compatible with developed speckles. The flow in the right vessel was twice as fast as the flow in the left one, and the simulations were repeated with different distances between the blood vessels, normalized by the width of the PSF. The second type of numerical simulation emulated blood flow through a bifurcation displaying the ability of the proposed method to reconstruct different vascular geometries. Spatiotemporal decompositions were not applied in any of these numerical simulation experiments, enabling us to test the super-resolution capabilities of sparse representation within each Doppler band.

Next, we evaluated the expected effects of the Doppler slicing and PSF estimation steps on the sparse representation results. To do so, we tested the performance of the sparse reconstruction algorithm on noisy data and performed reconstructions with distorted PSF estimations. Different levels of additive noise were used, resulting in SNR values between 0 and 100 dB, defined as the ratio between the maximum value of the simulated vessel and the standard deviation of the noise. The SNR level of the autocorrelation images was measured to be 10.5 dB higher than the original simulated signal; these values are used for reporting the results of the sparse representation algorithm. The dynamic range used for the display of Doppler slicing images was calculated as 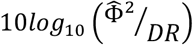 as typically done when presenting power Doppler images. However, in all other cases, we use the regular definition of 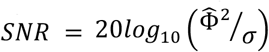, where *σ* is the standard deviation of the noise, to better fit previous sparse representation papers. To test the robustness of the proposed algorithm to PSF estimation inaccuracies, the PSF used in solving the inverse problem was scaled in the lateral direction to 75-125% of the original PSF. Together, these numerical simulations determine the desired PSF estimation precision and the SNR needed for reliable blood vessel separation following Doppler slicing.

### B. In vivo experiments

To test the proposed method experimentally, we used mouse brain scans. The animals underwent a craniotomy creating an acoustic window over the region of interest. All the scans were acquired using a Ventage 128 ultrasound system (Verasonics, Kirkland, WA, USA) and a Domino linear probe (Vermon, Tours, France) with a central frequency of 15 MHz (pitch: 0.11 mm). We acquired scans with 1000 Doppler ensembles. Each ensemble contained 200 pulse-echo acquisitions and lasted 0.2 s (frame rate of 1 kHz). Each pulse-echo acquisition was composed of 13 transmission angles (−6° to 6°) compounded together. The RF data was beamformed on a *λ*/2 spaced grid using Verasonics’ proprietary beamforming algorithm, and the tissue clutter was removed using SVD filtering [25]. The scan was then automatically separated into 1.6 × 1.6 mm patches with 75% overlap in each direction. All the processed patches and bands were summed to produce the final images, except for the ten bands with the slowest flow in each direction, due to their lower SNR level.

Following each Doppler scan, we performed a super-localization scan of the same imaging plane. These additional scans served as ground truth for testing the non-contrast super-resolution reconstructions. Definity (Lantheus Medical Imaging Inc., N. Billerica, MA, USA), a clinically approved contrast agent, was injected into the tail vein of the mouse at a concentration of 40 μL/Kg. Three microbubble injections were performed for each imaging plane. To separate single bubbles for ULM processing, the first 30 and last 60 SVD components were removed [30]. The PSF of the system was estimated by averaging over the envelope of single bubble signals, as in [10].

It was estimated once and then used for processing the non-contrast ultrasound scans. All the protocols were approved by Caltech’s Research Institutional Review Board.

## VI. Results

### A. Doppler Slicing Produces Clear Decompositions of Vascular Ultrasound Scans

We first studied the ability of Doppler slicing to decompose vascular ultrasound scans and investigated how the quality of the resulting decomposition varies with the number of ensembles. Classic power Doppler without contrast agents and directional power Doppler scans (**Fig. 2a-c**) were used as a reference, showing the fidelity and quality of the Doppler slicing results. The speckle noise in each of the Doppler bands was reduced by combining the information over many ensembles (**Fig. 2d** vs. **Fig. 2e** and **Fig. 2f**). Smaller vessels appear in the middle of the Doppler spectrum, and larger blood vessels inhabit the ends of the spectrum (**Fig. 2f** and **Supplementary Movie 1**). The reduction in speckle-noise level was quantified by calculating the structural similarity (SSIM) between Doppler bands composed of different numbers of ensembles and the reference Doppler bands calculated with 1000 ensembles (**Fig. 2g**). The structural similarity levels increase with the number of ensembles and are above 0.99 when 256 ensembles are used. Overall, Doppler slicing results in a clean hierarchical decomposition of the vasculature into a series of sparser bands.

**Figure 2:**
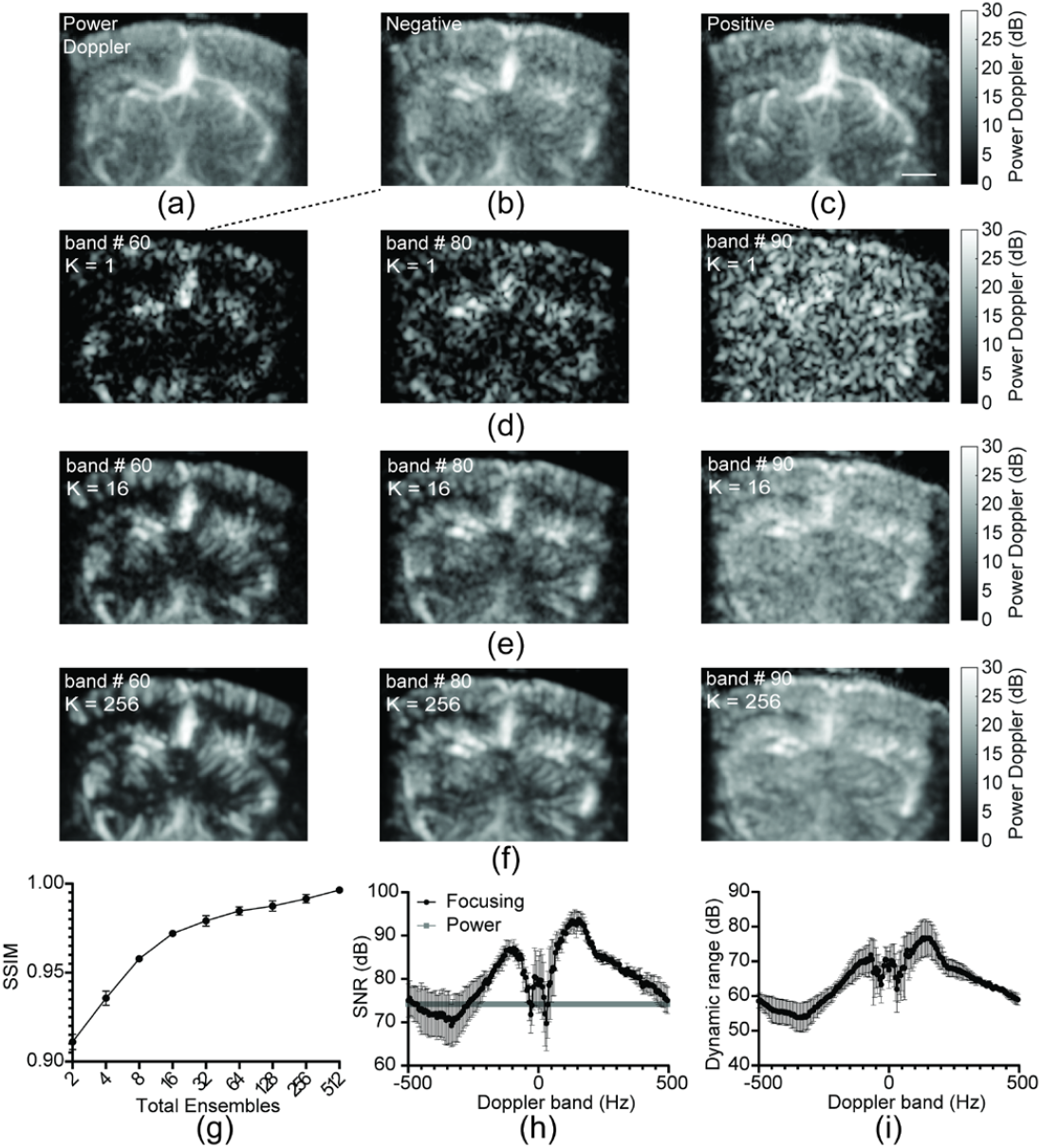
Doppler slicing produces sparser vascular images with high SNR and dynamic range. a. Standard power Doppler image of a mouse brain. b. Negative directional Doppler image of the same imaging plane. Decomposition to upward and downward flow separates close by vessels with opposite flow directions. c. Positive directional Doppler image. d. A few single-ensemble Doppler bands showing dominant speckle-noise patterns. e. Doppler bands calculated from 16 ensembles demonstrating improved vessel depiction. f. Doppler bands computed from 256 ensembles displaying clean maps of the different vascular structures. g. The structural similarity between the Doppler slicing bands and the reference images calculated from the entire dataset (1000 ensembles) increases with the number of integrated ensembles (n=5 scans). h. Improvement in SNR in low-frequency Doppler bands that contain microvascular structures (n=3 scans). i. High dynamic range is maintained over the entire range of Doppler slicing bands (n=3 scans). Bar plots show mean ± SEM (g-i). Scale bar is 1 mm (c).

The cranial vascular scans were used for evaluating two additional image quality parameters: the level of additive thermal noise and the dynamic range of the bands. A region of interest located outside the mouse brain was used for estimating the thermal noise level in each Doppler band (**Supplementary Fig. S1**). Doppler slicing spreads the additive thermal noise over all the bands, resulting in high SNR levels in the middle of the frequency range and a clear depiction of the microvasculature (**Fig. 2h**, and **Fig. 2f**, right). At the same time, the proposed approach virtually removed the stronger signal of large vessels (**Fig. 2f**, left) from the microvasculature bands (**Fig. 2f**, right), resulting in a high dynamic range throughout the bands (**Fig. 2i**). As we will see in the following sections, these sparser Doppler slicing bands and their high SNR levels and dynamic range lend themselves to better super-resolution processing.

### B. Simulations Show Sub-Diffraction Reconstructions Under Conditions Compatible with Doppler Slicing

After studying the ability of Doppler slicing to separate different vascular structures and quantifying its characteristic SNR and DR levels, we examined how these conditions could translate to vascular super-resolution imaging via numerical simulations. We tested the separation of close-by vessels under ideal SNR conditions and PSF estimations and then evaluated the robustness of the sparsity-based reconstruction to different noise levels and errors in PSF estimations.

The simulated Doppler data included a large number of scatterers per resolution cell with random amplitudes, resulting in developed speckles (**Fig. 3a**). Sparsity-based reconstruction was able to resolve vessels that were fused in the diffraction-limited and autocorrelation images (**Fig. 3b-d**). Cross-sections through vessels with different separations showed improved delineation (**Fig. 3e**, left) and resolution (**Fig. 3e**, right). Full-width-at-half-maximum (FWHM) measurements showed a significant improvement in vessel delineation of separable vessels. The FWHM of the sparse representation estimation was thinner by a factor of 3.87 compared to the autocorrelation signal and by a factor of 5.82 compared to the PSF. Looking at the full range of vessel spacing, we also observed a significant improvement in target separation (Rayleigh Criterion): a 2.28-fold improvement in vessel separation compared to the autocorrelation signal, 2.54-fold improvement compared to the temporal mean, and 3.13-fold improvement compared to the diffraction limit. These results demonstrated the expected contribution of Doppler slicing, as vessels that are separated into different Doppler bands will be better resolved in the final vascular map. At the same time, a simulated bifurcation was successfully estimated, demonstrating the ability of the proposed method to reconstruct arbitrary vascular structures without any shape priors, even if they reside in the same Doppler band (Supplementary Fig S2).

**Figure 3:**
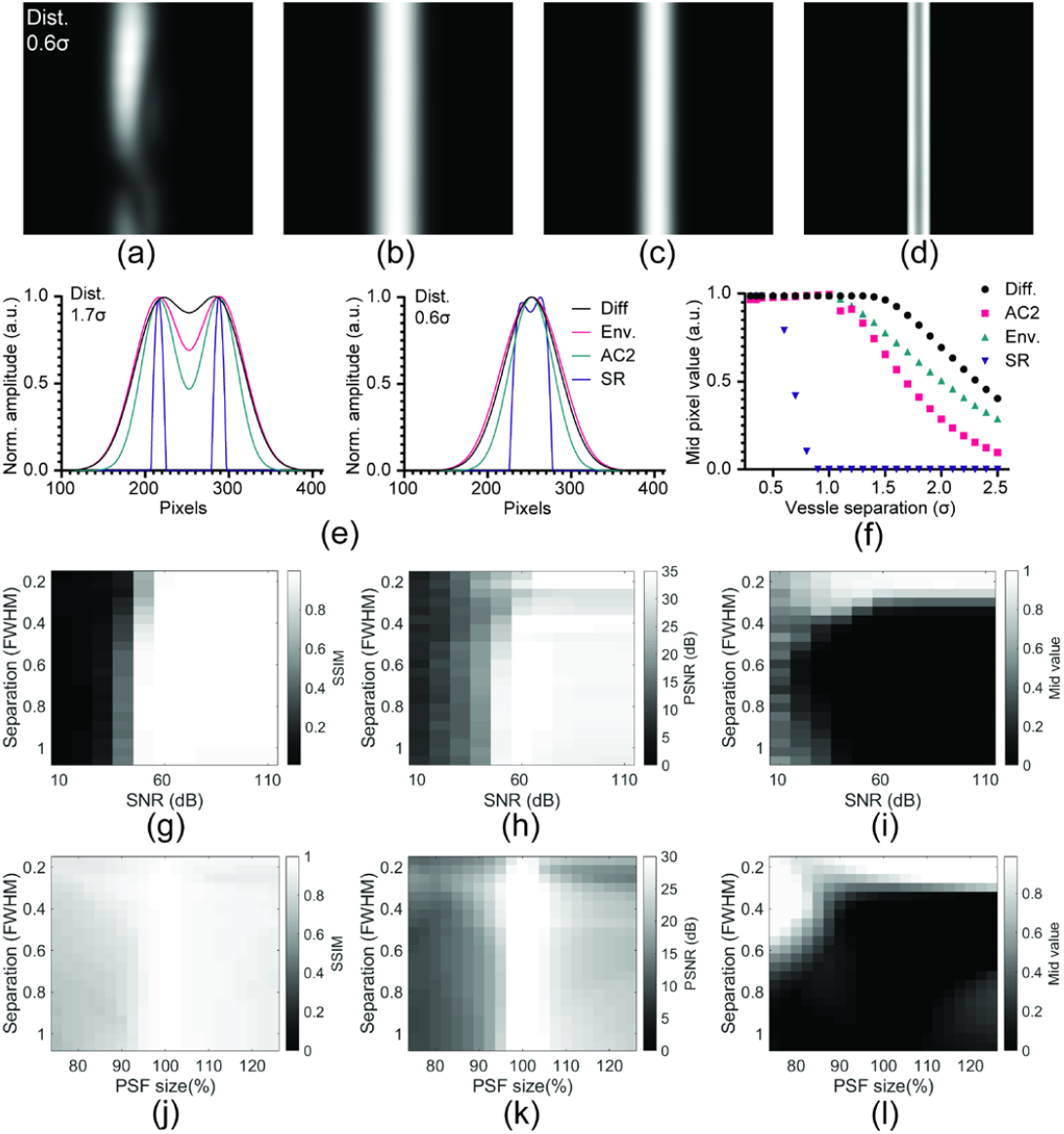
Numerical simulations demonstrate flow-line separation under conditions compatible with Doppler slicing. a. Single frame from simulation of flow inside two parallel vessels. **b**. Diffraction-limited temporal averaging cannot separate between the two close by vessels. **c**. Autocorrelation (power Doppler) image presents a tighter flow profile without resolving the two close by vessels. **d**, Sparse representation produces a clear reconstruction of the two neighboring flowlines. **e**. Cross-section from numerical simulation of far blood vessels presents improved FWHM (left, **d** = **1. 7 σ**) while cross-section from a simulation of close blood vessels shows improved vessel separation (right, **d** = **0. 6 σ**). **f**. The mid-pixel level achieved by the different processing methods is displayed over the full range of vessel distances, revealing improved resolution using sparse representation reconstruction. **g**. Structural similarity measurements show robust vessel reconstruction in the SNR levels characteristic of Doppler slicing. **h**. PSNR measurements present successful vessel reconstruction over the relevant noise levels. **i**. The mid-pixel value map shows stable vessel separation over this SNR range. **j, k**. Structural similarity and PSNR values (respectively) drop as the width of the PSF used for reconstruction deviates from the one used in the simulations. **l**. Blood vessel separation decreases when thin PSF estimations are used for reconstruction, while wide PSF estimations result in artifacts, even for highly separated vessels. Bar plot shows mean ± std (f).

Contaminating the simulations with different levels of additive noise helped determine the desired signal-to-noise level for reliable super-resolution reconstruction. We used structural similarity and peak SNR measurements to quantify the reconstruction quality for each SNR value and vessel distance. Both matrices showed good performance above SNR = 50dB with SSIM values above 0.97 and peak SNR values higher than 25.5 dB. Bellow SNR = 30dB (**Fig. 3g, h**), we observe problematic reconstruction with SSIM values dropping below 0.05 and peak SNR values lower than 12 dB. A deeper insight into this transition range can be obtained by looking at the pixel values in the middle of each cross-section as a function of the SNR and vessel distance (**Fig. 3i**). As the SNR decreases below 50dB, so does the vessel separation. At an SNR of 30 dB and below, we start seeing reconstruction artifacts even for highly separated vessels in the form of high mid-pixel value. The artifact level increases with lower SNR values. These results, showing robust reconstruction above SNR = 50dB, comply with the SNR levels produced by the Doppler slicing decomposition (**Fig. 2h**).

Finally, we tested the sensitivity of the reconstruction algorithm to errors in PSF estimation. Both structural similarity and peak SNR dropped for PSF estimations that were significantly wider or narrower than the kernel used in the simulation (**Fig. 3j, k**). SSIM values drop below 0.9 for PSF width below 95% or above 115%, while peak SNR values drop below 17.5 dB outside this range. Looking at the mid-pixel value matrix, we can observe that narrow PSF estimations resulted in reduced vessel separation, and wide PSF estimations showed artifacts even for highly separated vessels (**Fig. 3l**). These results demonstrate how important is accurate PSF estimations for successful sparse representation of non-contrast Doppler data.

### C. In-vivo Super-resolution Vascular Imaging using Doppler Slicing

Finally, we tested the ability of Doppler slicing to facilitate super-resolution imaging of dense vascular scans of mouse brains. Looking at a 1.6 × 1.6 mm image patch taken from the mouse cortex, we can see the vascular decomposition achieved by Doppler slicing, compared to the original power Doppler image (**Fig. 4a-c**, vs. **Fig. 4d**, respectively). Sparse representation improves the delineation of details that are blurred in the autocorrelation image of each band (**Fig. 4e-g** vs.

**Figure 4:**
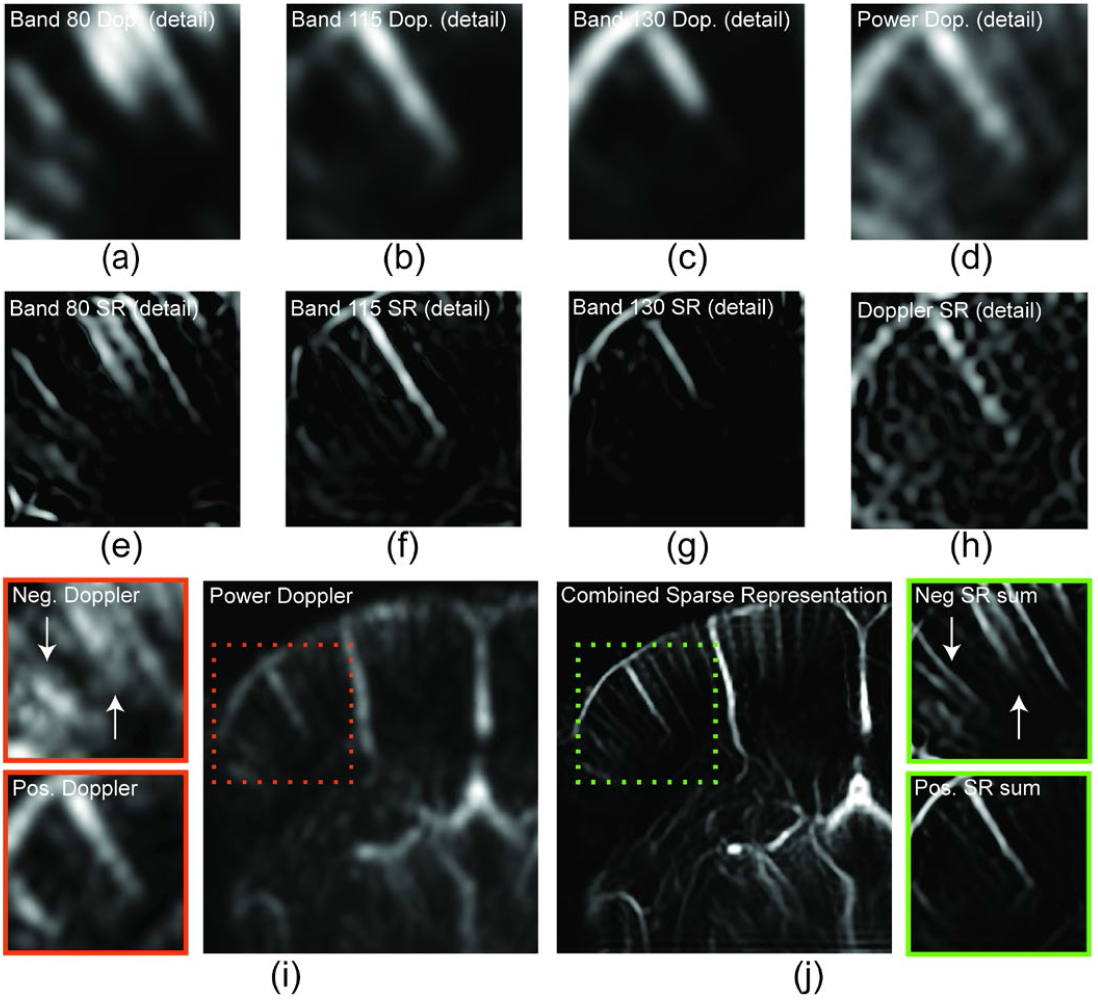
Doppler slicing facilitates super-resolution vascular imaging without contrast agents. a. An image patch from a negative Doppler band showing small vessels in the mouse cortex. b. An image patch from a positive Doppler band showing small vessels different from those detected in (a). c. An image patch from a different positive Doppler band showing mid-size vessels. Doppler slicing separates vessels of different sizes and flow directions. d. Standard vascular map of the same image patch. e-g. Sparsity-based super-resolution images of the Doppler bands in (a-c), respectively. h. Sparse representation of the power Doppler patch in (d). The high noise level and dense vascular structures in the power Doppler image impede the use of sparse representation processing. i. Power Doppler image of the right brain hemisphere (right). Negative and positive single patch Doppler images display low-resolution vascular details (left, top, and bottom, respectively) j. Summing the sparse representation reconstructions of all the Doppler bands. The resulting negative (right, top) and positive (right, bottom) frequency patches show vascular details that could not be resolved in the original directional Doppler images (panel i, left, top, and bottom, respectively). Combining all the Doppler bands and patches provides a super-resolved vascular map of the right brain hemisphere (left). Image patches are 1.6 × 1.6 mm (a-i).

**Fig. 4a-c**, respectively). Such improvement was not observed when applying sparse recovery to the original power Doppler image (**Fig. 4h**). The dense and noisy power Doppler signal (**Fig. 4d**) resulted in distorted reconstruction (**Fig. 4h**).

Combining all the processed bands, we can see how the sparsity-based reconstruction of the decomposed data revealed vascular information that wasn’t observable in the original directional Doppler data (arrows, **Fig. 4i**, left vs. **Fig. 4j**, right). Combining all the patches of different areas of the mouse brain, we observe a full image of the left hemisphere of the mouse brain. The processing of all the bands and patches was performed using the same set of parameters, showing that the proposed method does not suffer from overfitting and that it can be generalized to full scans.

Super-localization microscopy was used as ground truth for evaluating the result of the proposed algorithm. Scans of the same image planes were acquired with and without contrast agent injections. The sparse recovery image is highly compatible with super-localization microscopy, revealing vascular information that was masked in the power Doppler images (**Fig. 5a-c**). Cross-sections through several vessels show the improved separation of blood vessels using the proposed method in the lateral and axial directions (**Fig. 5d, e**, respectively). Dashed lines in the power Doppler image (**Fig. 5a**) mark the locations of these cross-sections. Cross-sections through small vessels (**Supplementary Fig. S3a, b**) were used to quantify the improvement in spatial resolution achieved by the proposed method. The average FWHM of these vessels was 42μm (**Fig. 5f**), a 2.4-fold improvement compared to the wavelength and close to 4-fold improvement compared to the FSF. These results demonstrate the potential of performing super-resolution imaging of vascular ultrasound scans of blood flow while showing that additional improvement in resolution can be gained when the use of contrast agents is optional.

**Figure 5:**
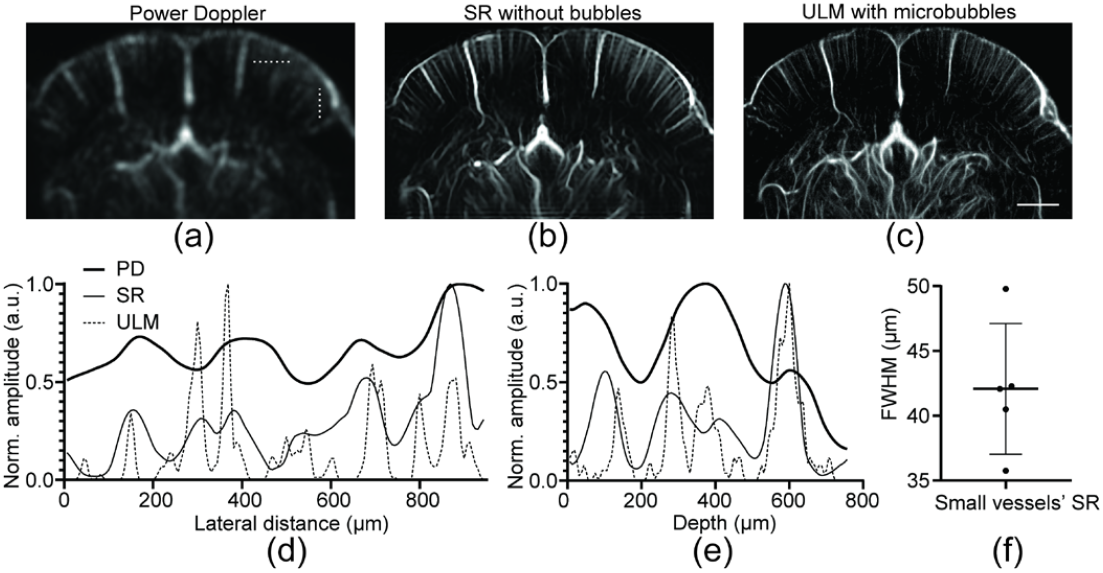
Sparse representation of Doppler bands shows high agreement with ultrasound localization microscopy. **a**. Standard power Doppler map of a mouse brain. **b**. Sparsity-based super-resolution image with Doppler Slicing revealing additional details that could not be resolved using power Doppler. **c**. Matching super-localization microscopy image showing close agreement with the sparse representation image in (b). **d, e** Cross-sections through small vessels show improved blood vessels’ separation using sparse representation in the lateral and axial directions, respectively. Dashed lines in (a) mark the locations of these cross-sections. **f**. FWHM measurements of cross-sections through small, isolated vessels show sub-diffraction resolution in sparse representation reconstruction. Scale bar is 1 mm (c)

## VII. Discussion

This work presents a framework for vascular ultrasound super-resolution imaging without contrast agents. We show that, in some instances, dense and overlapping blood vessels can be separated without microbubble injections. To do so, we integrated the vascular information provided by Fourier transforms over several ensembles, thus extending the processing beyond single power Doppler acquisitions. This approach resulted in a high SNR hierarchical decomposition of the vasculature that facilitates vascular super-resolution imaging by exploiting sparsity. The information needed for this reconstruction, including the PSF, can be estimated from the data. Even though this work looked at an iterative super-resolution implementation, Doppler slicing could be integrated with other computational super-resolution approaches such as iterative algorithm unfolding or deep learning methods [13], [14].

Doppler slicing and subsequent non-contrast super-resolution imaging could have several potential applications beyond pre-clinical research. For example, such capability could be used for imaging of changes in the morphology of microvasculature following anti-cancer treatments such as anti-angiogenic drugs. Tumor vessels tend to be tortuous and unorganized [34], [35]. It was shown that the success of specific anti-cancer treatments could be evaluated according to their effects on the morphology of the microvasculature that undergoes vascular ‘normalization’ [34]. Many of these changes occur at vascular scales below the resolution of classic ultrasound imaging methods. Another scenario in which the structure of the microvasculature is known to have clinical importance is the imaging of the vasa vasorum inside arteriosclerotic plaques [36]. Hyper vascular plaques tend to be more fragile, resulting in an increased risk of stroke. By improving the spatial resolution of vascular scans without using contrast agents, this technique could help to bring next-generation ultrasound imaging to underserved populations and remote communities.

Despite the capabilities presented in this work, this approach has several limitations. Contrast-enhanced ultrasound scans produce stronger vascular signals that enable the detection of capillary flow [17] and improve the imaging of blood flow through intact skulls [3], [5], [37], [38]. In addition, exploiting bubble sparsity resulted in more than a ten-fold improvement in spatial resolution. Thus, super-localization methods with contrast agent injections can detect and resolve smaller vessels. Currently, an acquisition time of around three minutes is needed to generate a contrast-free super-resolved vascular image using Doppler slicing, similar to the duration of plane wave super-localization scans [1]. Reducing the number of ensembles needed for Doppler slicing processing would save expensive scan time and reduce the effects of motion artifacts. Therefore, use of more advanced denoising methods for the removal of speckle noise from the single-ensemble Doppler slicing measurements is a subject of continuing work. Another line of research is extending the use of Doppler slicing to machine learning-based super-resolution methods. These approaches enable a more straightforward estimation of the PSF in different locations and tissues. In plane-wave ultrasound scans, the PSF changes according to the target’s location relative to the transducer. Assuming that the PSF is piece-wise constant, many estimations can be performed and matched to the processed image patches. Together, the improved SNR and precise PSF estimation enabled by these approaches have the potential to facilitate the detection and separation of smaller vessels.

## Supporting information

Doppler_slicing - supplementary movie 1

Doppler_slicing - SI

