## Supplementary material for "Doppler Slicing for Ultrasound Super-Resolution Without Contrast Agents": Doppler_slicing - SI

**Contents:**

Supplementary Figures 1-3

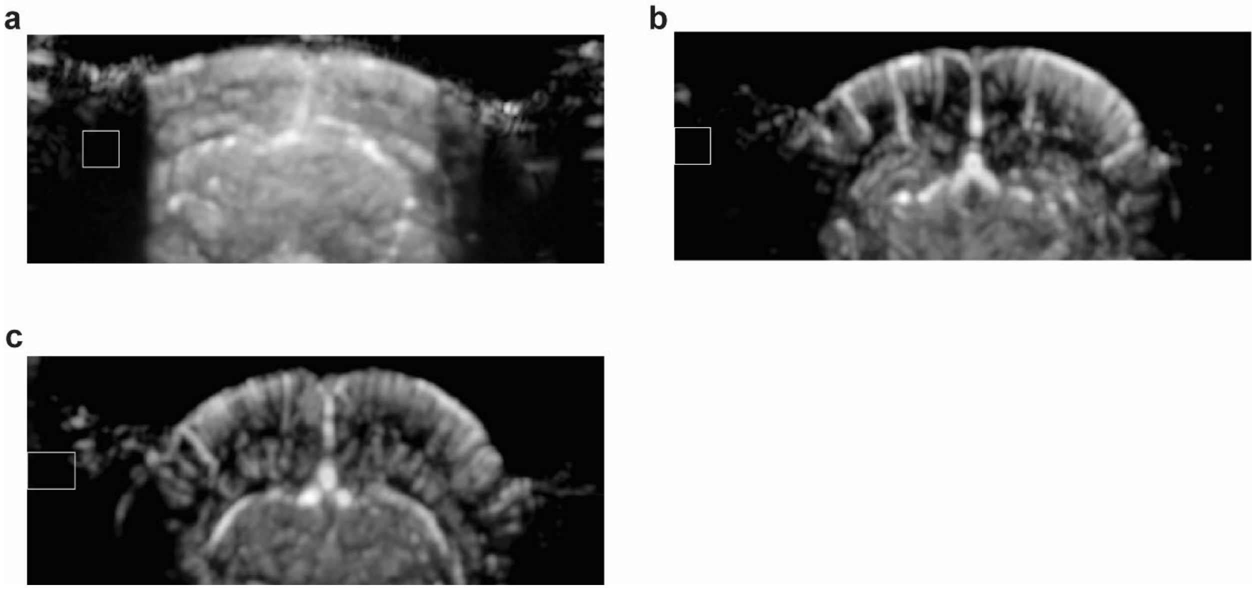


Supplementary Figure S1: **SNR estimation: ROIs used for noise level estimation. a-c**. ROIs used for calculation of the noise level in each scan as part of Fig. 2h, i.

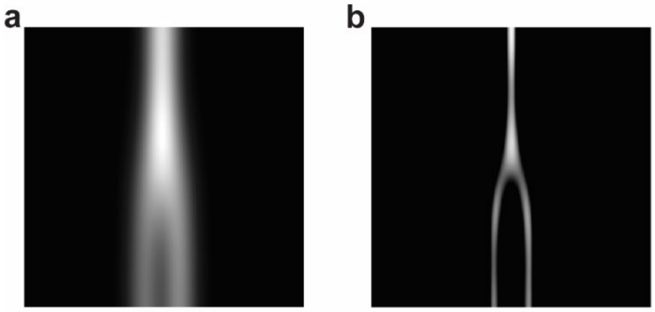


Supplementary Figure S2: **Additional simulation result: reconstruction of a bifurcation. a**. Autocorrelation (Doppler) image of the simulated bifurcation. **b**. Super-resolved version of the simulated bifurcation showing reconstruction of the bifurcation geometry.

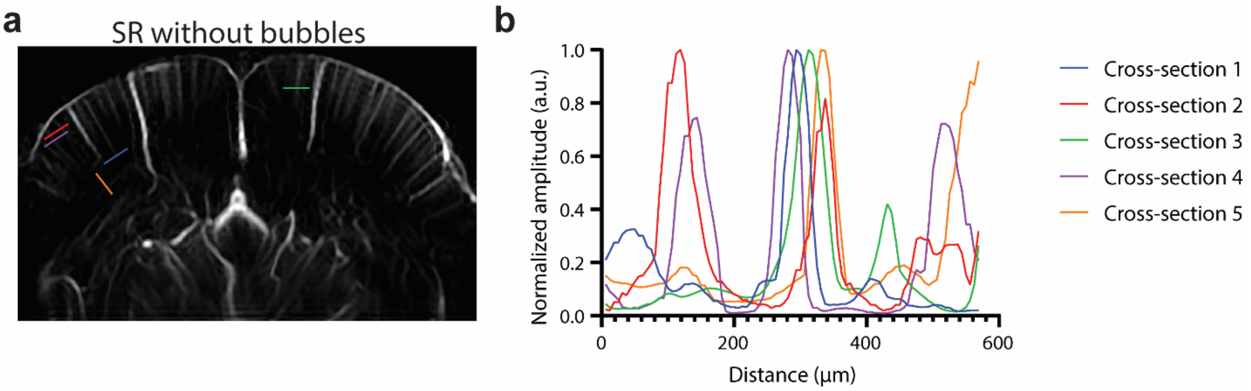


Supplementary Figure S3: **Small vessel FWHM: cross-sections. a.** Locations of cross-sections used for calculating the FWHM of small vessels (presented in Fig. 5f). **b.** Cross-sections used for calculating the FWHM of small vessels.
